# Dorsal Hippocampal Activation Suppresses Neuropathic Pain Behaviors: Chronic pain as extinction-resistant pain-related memory traces

**DOI:** 10.1101/292094

**Authors:** Wei Xuhong, W. Ren, M.V. Centeno, D. Procissi, Ting Xu, R. Jabakhanji, M. Martina, J. Radulovic, D. J. Surmeier, X.G. Liu, A.V. Apkarian

**Author notes:** Corresponding author: A Vania Apkarian.

## Abstract

Accumulating evidence suggests the hippocampus being involved in, and modified with, chronic neuropathic pain. However, it is still not clear whether hippocampal activity has direct control over neuropathic behaviors. Here we show that activation of the dorsal, but not ventral, hippocampus, by glutamate microinjection or by chemogenetically increasing excitability (PSAM/PSEM), completely or partially reversed neuropathic behaviors: tactile allodynia and thermal hyperalgesia in the models of spared nerve injury and lumbar spinal nerve ligation. Using a new methodology (chemo-fMRI), where we combine awake resting state brain imaging with viral vector mediated chemogenetic activation (PSAM/PSEM), we could demonstrate that increased excitability of dorsal hippocampus neurons altered resting state functional connectivity within circuitry specifically related to the extent of diminution of neuropathic behavior (tactile allodynia). The identified circuitry most reliably (survived a validation procedure) identified dorsal hippocampal connections to the somatosensory cortex and the thalamus. Moreover, anterograde tracing indicated non-overlapping projections from dorsal and ventral hippocampus. Thus, the present study exhibits a novel causal role for the dorsal hippocampus, and mediating circuitry, controlling neuropathic pain-related behaviors. Altogether, these results imply downregulation of dorsal hippocampus circuitry in chronic neuropathic pain; the activation of which reverses pain behaviors either through disruption of accumulated memories and/or by enhancing extinction circuitry.

## Introduction

Accumulating evidence implicates hippocampal circuitry in chronic pain. Animal and human studies now show hippocampal processes being engaged in the transition to chronic pain, and multiple hippocampal mechanisms are now documented to be altered in the chronic pain state.

Rodent models for chronic pain, where partial peripheral nerve injury commonly leads to persistent pain-like behaviors, are now shown to concomitantly lead to a multiplicity of hippocampal dysfunctions, including altered synaptic plasticity, abnormal cytokine expression, decreased expression of NMDA receptor subtypes, instability of place cells, and a reorganization of resting state connectivity [5; 9; 10; 27; 32; 38; 54]. Moreover, adult hippocampal neurogenesis seems to be a critical component of the mechanism controlling the transition to chronic pain [3], and in turn rate of hippocampal neurogenesis is shown to diminish once chronic pain is established [32]. Consistent and complimentary evidence is now also observed in humans. Multiple somatic chronic pain conditions exhibit hippocampal volumes smaller than in matched healthy controls, as reported for chronic back pain and for chronic complex regional pain syndrome [32], and also for chronic knee osteoarthritis [49]. The hippocampal volume itself is a risk factor for subjects transitioning from sub-acute back pain into chronic back pain [49], and the transition to chronic back pain is accompanied with functional connectivity changes both intrinsically within the nucleus, and between the hippocampus and the cortex [31]. Thus, hippocampal physiology and connectivity undergo large adaptations with the establishment of chronic pain. Yet, how these changes influence on the ability of hippocampal circuitry to control pain behavior remains unclear, and this is the main question we address in the current study.

Ample evidence demonstrates that the hippocampus is involved at least in episodic memory, spatial cognition, and mood. Its role in chronic pain has been theorized based on observations of individual painful events giving rise to memory traces that could persist for a lifetime [2; 28], and thus chronic pain may be conceptualized as a state of continuous accumulation of conscious and/or sub-conscious memory traces that resist extinction. The hippocampus can be segregated functionally and structurally, in humans along the anterior-posterior axis, and in rodents in the dorsal and ventral axis [6; 45], differentiating between these divisions regarding genetic expression, cell type patterns, and efferent connectivity [19]. Functionally the dorsal division is primarily associated with spatial memories, contexts, and spatial navigation, and memory retrieval, consolidation, and in extinction processes [47]; while the ventral division is more involved in affective modulation especially regarding anxiety [1; 7; 19; 30; 35]. To our knowledge there is no information as to the specialization of these subdivision within the context of chronic pain. Here we examine the differential effects of modulating excitability of each sub-division on neuropathic pain behavior.

We hypothesize that hippocampal adaptations with the transition to chronic pain, specifically after a peripheral neuropathic injury in rodents, would enhance the involvement of hippocampal circuitry in pain behaviors. We test this hypothesis by studying the modulation of exaggerated tactile and thermal responses in rat models of neuropathic pain, following local activation of hippocampal neurons, either with microinjection of glutamate or by activating depolarizing conductances using chemogenetic viral vector manipulations. We also combined the chemogenetic manipulation with awake resting state-fMRI (chemo-fMRI) [13] to identify hippocampal circuit reorganizations underlying the modulation of neuropathic pain behaviors, and used anterograde tracing to identify the dorsal and ventral hippocampus projections.

## Results

### Microinjection of Glutamate into dorsal hippocampus, but not ventral hippocampus, transiently reverses chronic pain-like behaviors induced by neuropathic injuries

A series of experiments explored changes in pain-like behaviors in rats with neuropathic injuries. Peripheral SNI injuries, when performed in China, resulted in bilateral tactile allodynia (animals, 30%, where tactile allodynia was strictly ipsilateral were excluded from the study); while the same procedure when performed in the US induced tactile allodynia only for the hind paw ipsilateral to the injury, accompanied with essentially unaltered touch sensitivity for the contralateral hind paw. To differentiate these experiments we label them, Chinese-SNI (C-SNI) and US-SNI (U-SNI), respectively. Two groups of C-SNI neuropathic rats were tested for tactile allodynia, following unilateral (ipsilateral to the injury) dorsal hippocampal microinjections of either glutamate (21.2 pmol) or saline (**fig. 1 A**). Seven days after C-SNI, when bilateral mechanical allodynia was established (tactile thresholds decreased from a baseline of 25 g to about 1 g, 4 days after injury), glutamate, but not saline, reversed mechanical allodynia. Within 2 hours after injection of glutamate, but not saline, paw withdrawal thresholds on the hind paws ipsilateral and contralateral to the injury (and to the hippocampal injection) were already increased, and this reversal of tactile thresholds to near baseline levels lasted 2 days. Four days after the first injection of glutamate (11 days after C-SNI), when the effects of glutamate had subsided, a second injection of glutamate, but not saline, again similarly reversed tactile allodynia (see **Supplementary Table 1** for statistics).

**Fig. 1.**
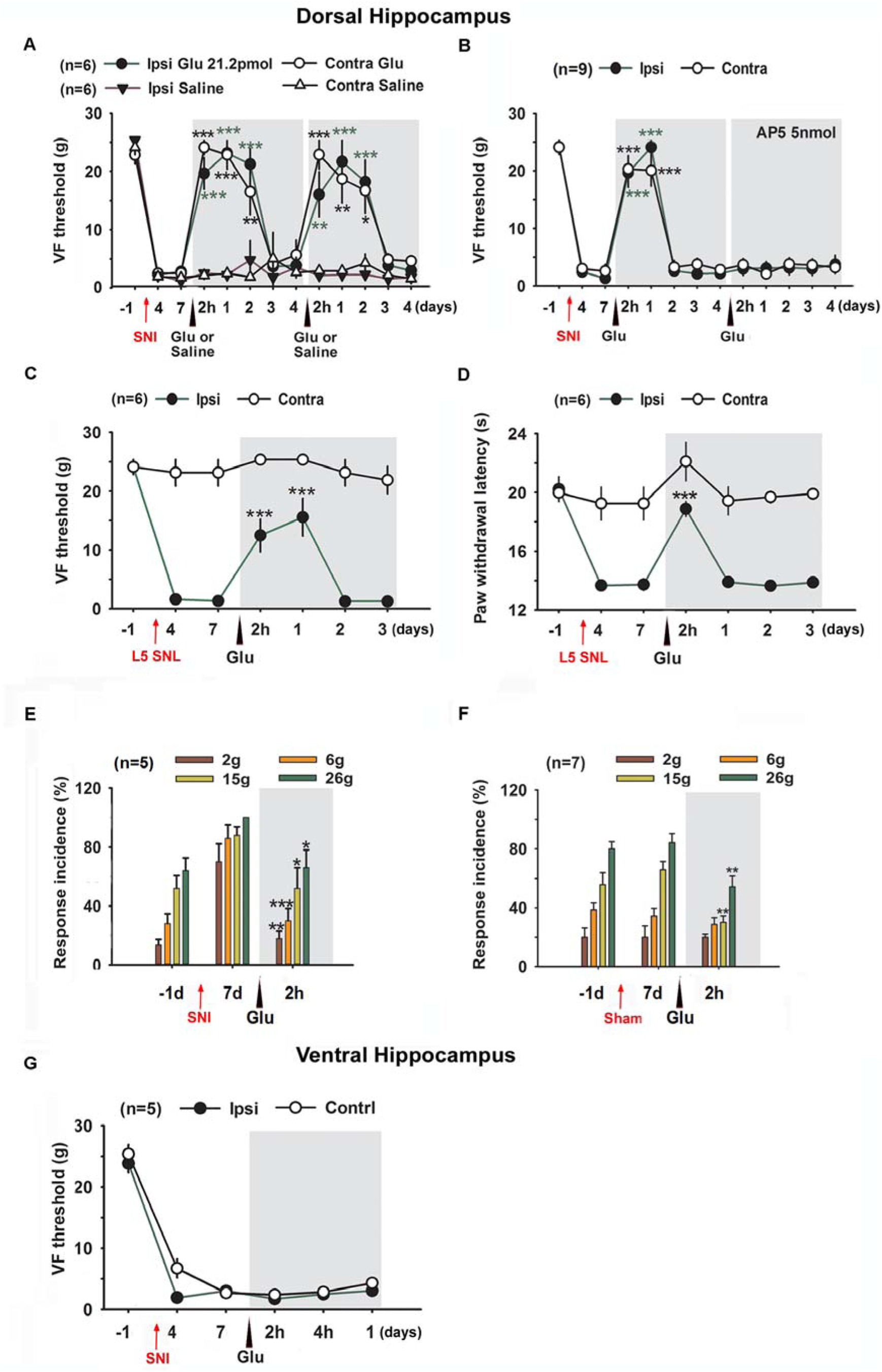
Glutamate microinjected into dorsal hippocampus, but not ventral hippocampus, alleviates chronic pain-like behaviors induced by neuropathic injuries. **A**. In two separate groups of Chinese-SNI (C-SNI, n=6/group), where peripheral nerve injury results in bilateral tactile allodynia, glutamate (21.2 pmol) but not saline, injected into dorsal hippocampus (ipsilateral to SNI injury) 7 days after SNI, reversed tactile allodynia on both ipsilateral and contralateral hind paws. Four days after the first glutamate injection (11 days after SNI), when the effect of original glutamate application had subsided, a second injection of glutamate, but not saline, again reversed tactile allodynia for both hind paws. In this and other panels, post-hoc statistical significance of differences from zero, from control groups, or from baseline conditions are indicated as follows: ***p<0.001, **p<0.01, *p<0.05, n.s. p>0.05, parametric or non-parametric tests, see **supplementary table 1** for all statistical tests. **B**. Dorsal hippocampal microinjection of glutamate at a lower concentration (5.3 pmol in 1µl volume) significantly increased the 50% paw withdrawal thresholds for both hind paws for 2 days. Four days after the first glutamate injection (11 days after SNI), when glutamate injection was preceded by an AP5 injection (through same cannula, 5nmol, 30 minutes before glutamate injection) the dorsal hippocampal glutamate failed to reverse mechanical allodynia, bilaterally. **C-D**. In the L5 SNL-induced neuropathic pain model, tactile allodynia (**C**) and thermal hyperalgesia (**D**) on ipsilateral hind paws were reversed with glutamate microinjected into dorsal hippocampus (ipsilateral to the peripheral injury, 5.3 pmol), with no effect on uninjured contralateral paw tactile and thermal sensitivities. **E-F.** The effect of glutamate injected into dorsal hippocampus on mechanical response incidence in two new groups of C-SNI (**E**) and Sham injury rats (**F**), for ipsilateral hind paw. Glutamate injection in dorsal hippocampus (5.3pmol, 7 days after SNI) in C-SNI reversed response incidences to pre-injury levels, across all forces; while in sham injured animals only responses to highest force were diminished. **G**. Microinjection of glutamate in the ventral hippocampus (5.3pmol, 7 days after SNI) had no effect on C-SNI-elicited mechanical allodynia, bilaterally.

In a second group of C-SNI rats we tested efficacy of a lower dose of glutamate (5.3 pmol), as well as the effect of pretreatment of glutamate with AP5 (**fig. 1B**). Lower dose glutamate injected into dorsal hippocampus, 7 days after SNI, could still reverse the mechanical allodynia on both hind paws, but the effect lasted for only 1 day. The effect of 5.3 pmol glutamate on mechanical allodynia was completely blocked with AP5 pretreatment (5 nmol, 30min before glutamate injection through the same cannula), tested at 11 days after SNI.

We next tested whether glutamate could affect pain behaviors on another neuropathic pain model: L5 SNL model; a model where the injury results in increased sensitivity for hind paw stimuli only on the injury side (study was done in China) (**fig. 1 C-D**). Both mechanical allodynia (**fig. 1C**) and thermal hyperalgesia (**fig. 1D**) for the ipsilateral, hypersensitive, hind paw were reversed by 5.3pmol glutamate injected unilaterally in the dorsal hippocampus (ipsilateral to the injury), whereas it did not perturb the contralateral hind paw tactile and thermal sensitivities, which were not changed by the L5 SNL injury.

In a new group of C-SNI and sham operated rats we also tested whether glutamate could affect the paw withdrawal incidences induced by four different forces of Von Frey hairs (2, 6, 15, 26g). In C-SNI animals, dorsal hippocampus microinjection of 5.3 pmol glutamate renormalized increased response incidences of the ipsilateral hind paw for all four Von Frey hair forces (**Fig. 1E**). However, in sham operated rats, the same dose of glutamate only decreased response incidences induced by the highest forces tested, 15g and 26g Von Frey hairs (**Fig. 1F**).

In contrast to the dorsal hippocampus, unilateral injection of glutamate into ventral hippocampus (5.3 pmol, ipsilateral to the injury) had no effect on C-SNI induced mechanical allodynia, bilaterally (**Fig. 1G**). Compared to pre-injection (day 7 post-SNI), there were no differences in bilateral paw withdrawal thresholds at 2-hours, 4-hours and 1-day after glutamate injection.

### Chemogenetic activation of dorsal hippocampal neurons reverses SNI-induced tactile allodynia

As glutamate is an excitatory transmitter and the glutamate effects could be abolished by blocking NMDA receptors, we hypothesized that glutamate may be modulating neuropathic pain by enhancing neuronal activity in dorsal hippocampus. To determine whether the activity of dorsal hippocampal neurons was responsible for tactile allodynia, dorsal hippocampal neurons were manipulated in vivo using a chemogenetic PSAM/PSEM approach [44]. The technique takes advantage of viral vectors to introduce excitatory 5HT3 channels in the cell membrane of neurons that replicate the virus. These channels are selectively activated by the ligand PSEM^89S^ and remain silent in its absence. Thus, neurons expressing the recombinant channel can be selectively activated in-vivo by i.p. administration of PSEM^89S^. To this end, the right and left dorsal hippocampus of wild type rats were injected with adeno-associated virus (AAV) carrying PSAM^L141F,Y115F^-5HT3 HC/green fluorescent protein (GFP) expression construct [44]. Immuno-histochemical analysis confirmed the expression of PSAM-5HT3-GFP in dorsal hippocampus (**fig. 2A**). In ex-vivo brain slices from infected rats, bath application of PSAM-5HT3 ligand -PSEM^89S^ - induced a reversible depolarization and generation of action potentials in GFP positive neurons (**fig. 2B, C,** n=6). Infected rats underwent SNI surgery following 4-week virus expression. These experiments were done in U-SNI rats (n=4), where injury increases response sensitivity primarily to the hind paw ipsilateral to the injury (SNI paw; from a baseline threshold of 30 g to lower than 10 g at 4 days after injury; **fig 2D**). Five days after U-SNI, rats received a single intraperitoneal injection of PSEM^89S^ (30 mg/kg). One hour after PSEM^89S^ administration, tactile allodynia of SNI paw was diminished (increased threshold) (**fig. 2 D, E**). In contrast, intraperitoneal saline injection (day 17 after SNI) had no effect on SNI paw tactile allodynia, in these PSAM-5HT3 infected rats (**fig. 2 D, F**). Nineteen days after SNI, we retreated these rats with PSAM^89S^, and a similar magnitude, transient, diminution of tactile allodynia was again observed for SNI paw, with a peak response at around 1 hour after administration of PSAM^89S^ (**fig. 2 D, G**). Meanwhile, neither PSEM^89S^ nor saline could modulate the tactile thresholds for the non-SNI paw, where tactile sensitivity was not different from pre-injury baseline (**fig. 2D, blue**). These results imply a causal relationship between excitability of dorsal hippocampal neurons and SNI-induced neuropathic tactile allodynia.

**Fig. 2.**
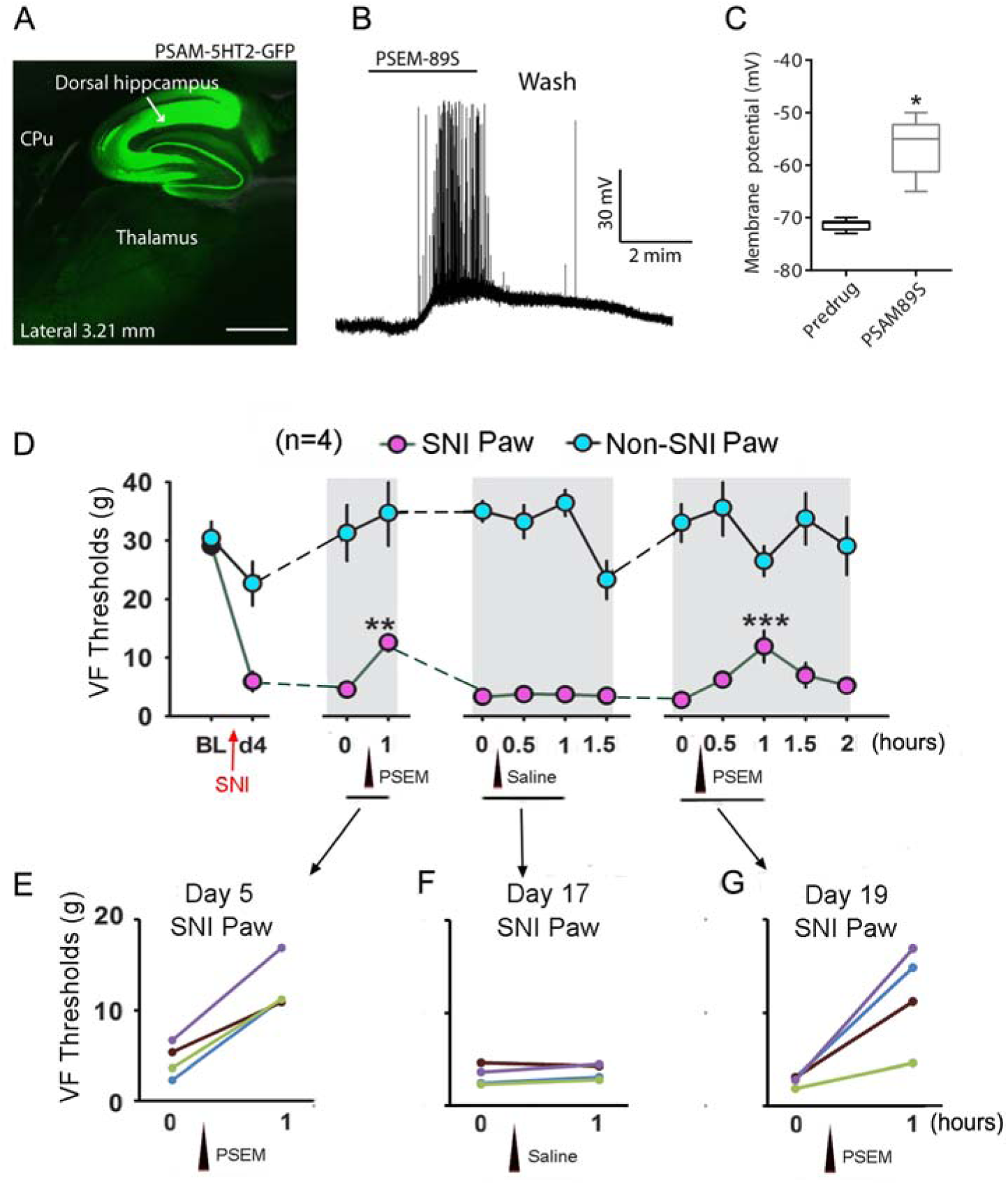
Chemogenetically increasing excitability of dorsal hippocampus neurons diminishes neuropathic tactile allodynia. **A**, In AAV-PSAM^L141F,Y115F^-5HT3 HC-GFP transduced rats, GFP fluorescence was exclusively expressed in dorsal hippocampal neurons (Scale bar = 11.5 mm). **B-C**, In the slice from infected mouse, bath application of PSEM^89S^ (10µM) elicited rapid depolarization and spiking in GFP-positive dorsal hippocampal neurons (n=6). **D-G**, In PSAM-5HT3 infected U-SNI animals (SNI injury only induces tactile allodynia on the SNI paw but not the non-SNI paw), intraperitoneal injection of PSEM^89S^ (30 mg/kg) rapidly and reversibly blunted tactile allodynia of SNI ipsilateral paw, on Day 5 and Day 19 after SNI, but had no effect on the contralateral uninjured paw (**D, E, G)**; whereas the treatment with saline did not perturb tactile sensitivity of SNI and non-SNI hind paws, on Day 17 after SNI (**D, F**).

### Chemogenetic activation of dorsal hippocampus changes functional connectivity, in proportion to reversal of tactile allodynia

To identify brain mechanisms controlling neuropathic pain with hippocampal activation, we combined chemogenetic manipulation of hippocampal neuronal excitability with awake, resting state fMRI scans, performed in awake trained U-SNI rats (chemo-fMRI). We specifically sought to unravel dorsal hippocampus functional connectivity changes during increased excitability of these neurons, in relation to neuropathic tactile allodynia changes.

Nine weeks prior to peripheral injury, wild type rats were injected with AAV carrying PSAM^L141F,Y115F^-5HT3-GFP in bilateral dorsal hippocampus (n=7). Two weeks later, they underwent head-post implantation (the only restraint used during brain imaging), and trained over 2-weeks for awake brain scanning [12; 13]. Peripheral unilateral SNI was then performed (U-SNI), a day after collecting baseline tactile sensitivity for both hind paws (BL VF). Four to five days after SNI: 1) tactile thresholds were measured; 2) brain scans were collected; 3) either PSAM^89S^ or saline was intraperitoneally administered (randomized), tactile sensitivity tested, and resting state fMRI collected twice; 4) about 2 hours later, the behavioral and brain scanning procedure was repeated, after administering the complementary intraperitoneal injection (those receiving PSAM^89S^ in the first round now received saline, and vice versa) (**fig. 3A**, experimental timeline). Four of the animals underwent this behavior testing and brain scan procedure on day 4 after SNI surgery; the rest underwent the same procedure on day 5.

**Fig. 3.**
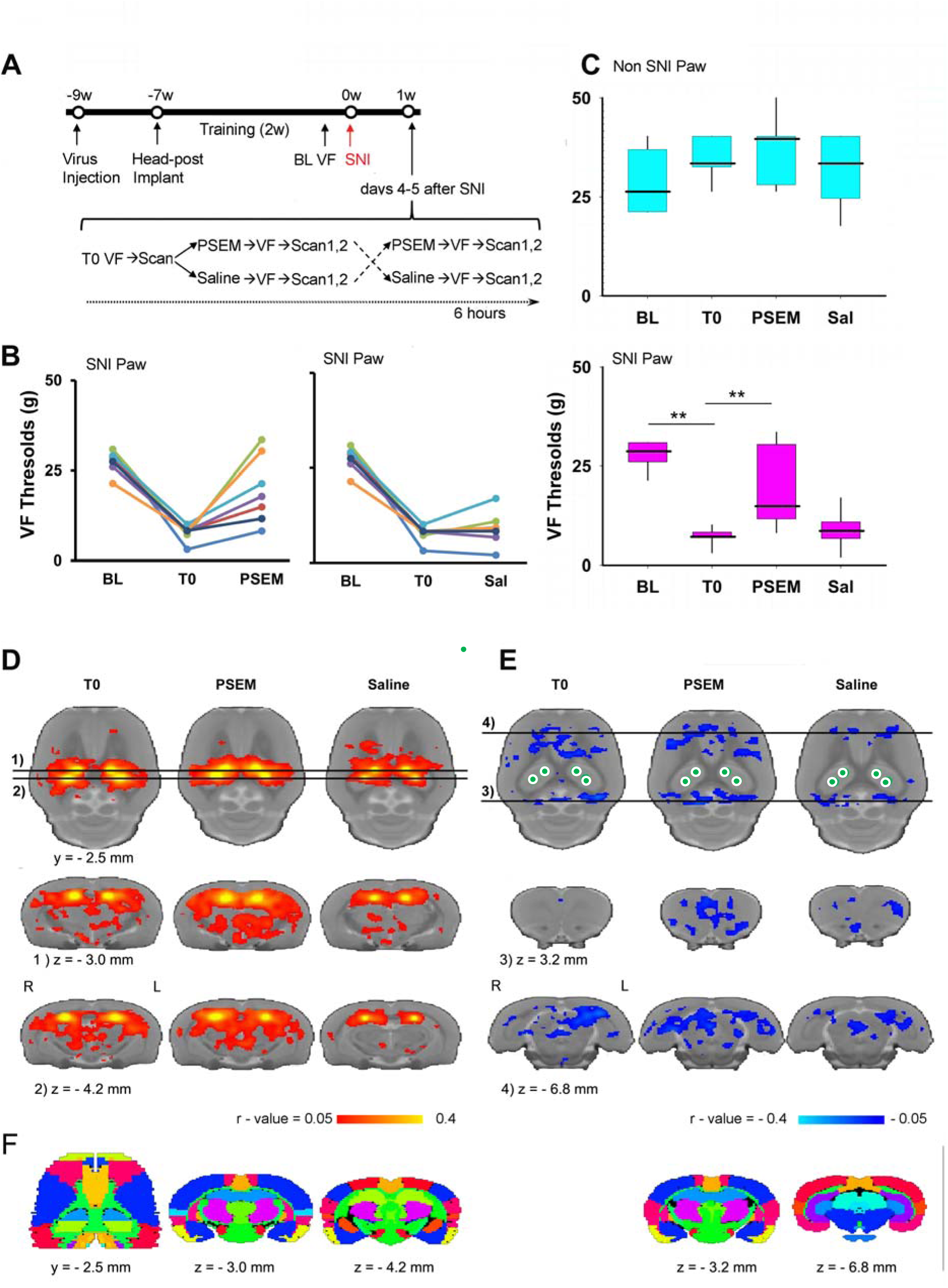
Awake chemo-resting-state-fMRI identifies dorsal hippocampal networks, with/without increased dorsal hippocampus neuronal excitability. **A.** Experimental timeline. Rats (n = 7) were first injected with AAV9-SYN-PSAM-L141F-Y115F-5HT3HC-GFP virus into bilateral dorsal hippocampus (~ 9 weeks prior to SNI surgery; U-SNI resulting in tactile allodynia limited to SNI hind paw). Two-weeks later they received implantation of a head-post (~ 7 weeks). These animals then underwent 2-week training period to enable awake resting state fMRI scanning. 1-2 days prior to SNI inju<u>ry, tactile thresho</u>lds were assessed for left and right hind paws (BL VF). SNI surgery was then performed unilaterally. 4 or 5 days after surgery, tactile thresholds (T0) were assessed and scanned for resting state fMRI, immediately afterwards animals received either saline or PSEM injections (i.p.), retested for tactile thresholds and again underwent awake resting state fMRI (2 consecutive scans). Two hours later, rats that had received PSEM were injected with saline, and vice versa, and tactile thresholds and fMRI were then assessed one more time. Scan 1 was used for discovering brain regions modulated with changes in excitability of dorsal hippocampus neurons; scan 2 data were reserved for validating outcomes. **B.** Individual animal tactile thresholds are shown for the SNI hind paw, at baseline (**BL**), 4-5 days after SNI (**T0**), and 1 hour after either **PSEM** or **saline** injection. In every animal, we observe increased tactile thresholds after PSEM but not after saline. **C.** Group median (quartiles and minimum/maximum) tactile thresholds are shown, for the non-SNI hind paw (blue) and SNI hind paw (pink). Tactile thresholds for the SNI hind paw diminished 1 hour after PSEM, but not after saline. **D.** Mean increased functional connectivity (FC) maps, averaged across all four seeds and animals. The three maps show mean FC, at T0: just prior to PSEM or saline injections; PSEM: about 1 hour after PSEM injection; Saline: about 1 hour after saline injection. **E**. Mean decreased FC maps, for the same conditions as in **D**. Top row: Brain maps at a single horizontal dorsoventral location. Bright yellow regions in D, and white circles in E, localize the four dorsal hippocampus seeds, where PSAM-5HT3 virus injected. Middle and bottom rows are coronal slices. Note: FC maps (in **D & E**) are intentionally un-thresholded to exhibit overall level of noise. **F.** Standard rat atlas slices corresponding to the FC map slices are displayed.

Consistent with the effects of PSEM^89s^ on U-SNI rats observed in the previous experiment, PSEM^89s^, but not saline, administration again reversed tactile allodynia of the SNI paw; and again, neither PSEM^89s^ nor saline administration affected non-SNI paw tactile sensitivity (**fig. 3B, C**). Thus, behaviorally the obtained results replicate those we observed in the animals studied in **figure 2**.

Group averaged (across all 4 dorsal hippocampus seeds, and all U-SNI animals) functional connectivity (FC) at 3 time points: 1) just prior to PSEM^89s^ or saline injections, T0; 2) for PSEM injection condition; and 3) for saline injection (using only scan 1 data for PSEM and saline conditions) were generated only for visual examination. Positive and negative average correlation coefficients are shown in **fig. 3 D and E**. It is visually apparent that the PSEM^89s^ condition shows more extended positive and negative FC (r-values) than T0 or saline conditions.

We next generated FC change (ΔFC) maps (between PSEM^89s^ and saline conditions; scan 1 data) and performed a correlation analysis to directly relate FC with behavior. Changes in tactile thresholds (ΔThreshold) for SNI paw were correlated with the FC contrast maps, between PSEM^89s^ and saline conditions, using the 4-seed average for each animal (scan 1 data). The resultant map indicates the brain regions where FC (to all 4 dorsal hippocampus seeds) changed in proportion to changes in tactile allodynia for the SNI paw, between PSEM^89s^ and saline conditions (**fig. 4**). Cluster corrected brain regions that relate FC and behavior (ΔFC with ΔThreshold) are shown in **figure 4A** (list of regions and statistical properties are in **supplementary table 2**). Brain regions with a positive relationship between FC change and touch sensitivity change (that is, areas where increased FC with PSEM^89s^ reflecting pain relief) were localized mainly to various cortices: primary somatosensory and motor cortices, multiple portions of the retrosplenial cortex, insular and medial prefrontal cortex; as well as dorsolateral thalamus, and parts of the dorsal hippocampus. On the other hand, brain regions where increased FC in PSEM^89s^ condition led to exacerbation of pain involved subcortical regions: zona incerta, dorsal and ventral pallidum, as well as parts of the primary somatosensory cortex and a region in the posterior hippocampus.

**Fig. 4.**
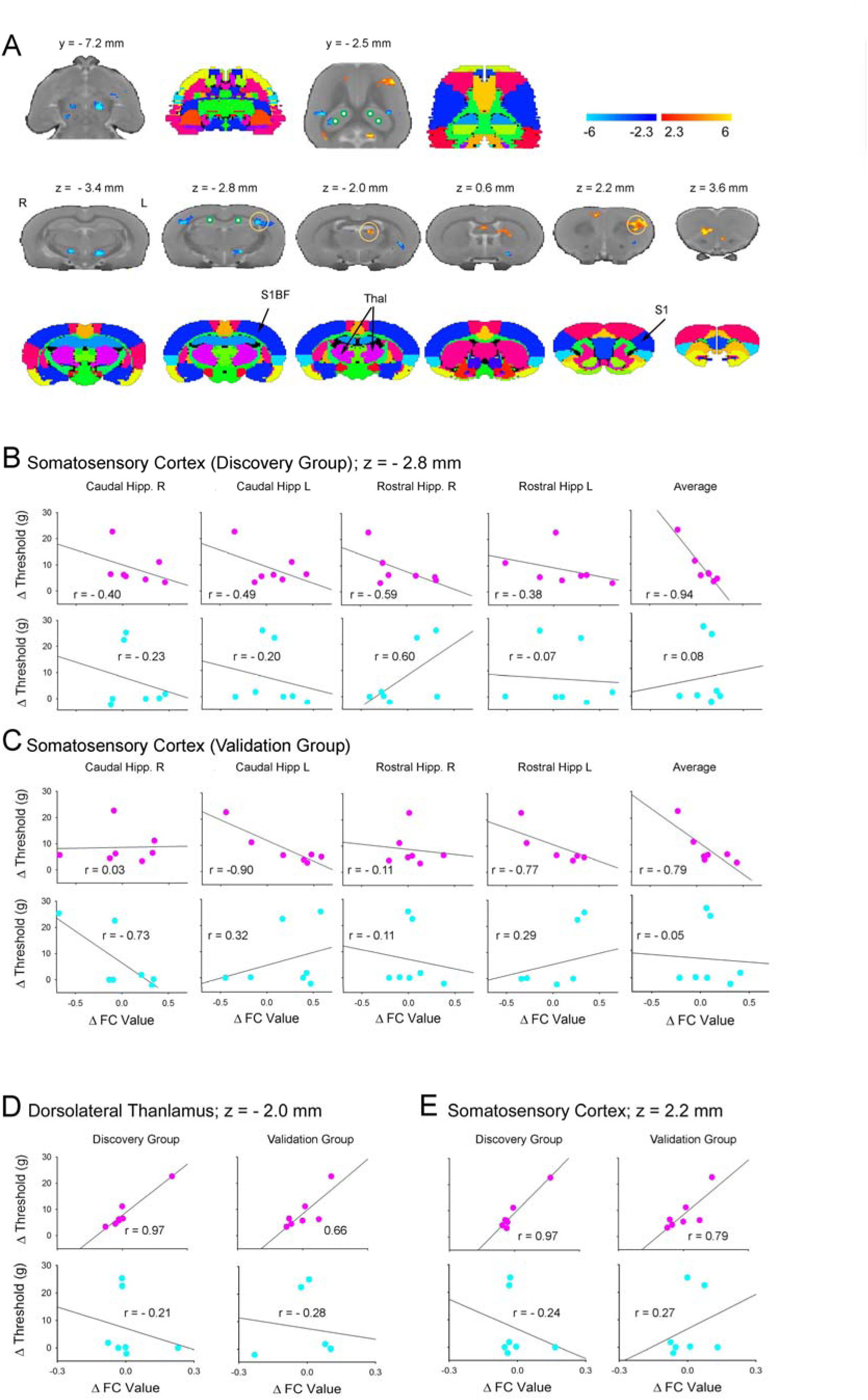
Identification of dorsal hippocampus driven functional networks that reflect changes in neuropathic tactile allodynia behavior as a function of changing dorsal hippocampal neuronal excitability. Using a rigorous discovery and validation approach, three dorsal hippocampus driven networks were identified, the properties of which were related to changes in tactile allodynia between PSEM^89S^ and saline conditions, specifically for the SNI hind paw. **A.** The map delineates brain regions where FC changes (ΔFC; between PSEM and saline conditions) were also correlated with change in tactile allodynia (ΔThreshold; between PSEM and saline conditions), for stimulating the SNI hind paw (using only scan 1 data) (cluster corrected for multiple comparisons; **supplementary table 1** summarizes identified regions and associated statistics). Standard atlas slices are also illustrated. **B.** An example brain region, primary somatosensory cortex (barrel field), where individual seed related ΔFC are shown relative to ΔThreshold (determined just prior to the brain scans), for both the SNI hind paw (top scatter graphs) and the non-SNI hind paw (bottom scatter plots). Right most panels are the average across all four hippocampal seeds, where we observe a strong relationship between ΔFC and ΔThreshold for the SNI hind paw but not for the non-SNI hind paw. **C.** The same relationship as in B, but for FC values extracted from the validation data (scan 2), using the coordinates derived from the discovery analysis. Again, we observe the SNI hind paw ΔThreshold is correlated with the average ΔFC, but not for the non-SNI hind paw. Thus, we consider this brain regional FC a validated outcome, indicating that high FC values in the region are related with high neuropathic pain. **D, & E.** Two additional brain regions identified in the discovery data could be validated. In both regions, discovery and validation results show a positive relationship between ΔFC and ΔThreshold for the SNI hind paw, but not for the non-SNI hind paw. Thus, in these 2 regions, high FC values (in PSEM^89S^) are associated with decreased neuropathic pain. Only across-seeds group averages are displayed.

For the top 8 highest-probability brain regions identified relating ΔFC with ΔThreshold, we extracted FC values from scan 2 data (using coordinates for each peak as identified in the scan 1 contrasts), for both PSEM^89s^ and saline conditions, and tested the relationship between resultant ΔFC to ΔThreshold (unbiased validation step). Only 3 brain regions survived this step (taking discovery and validation together, for SNI hind paw p<0.0025), and their properties are illustrated in **figure 4B-D**. **Figure 4B, C** shows discovery and validation scatter plots, for every hippocampal seed, for SNI and non-SNI hind paws, as well as the average scatters across all four seeds. Within this region, primary somatosensory cortex barrel field, correlations between ΔFC and ΔThreshold for SNI hind paw is quite variable for specific seeds, and generally weaker in the validation data; yet on average both discovery and validation r-values are statistically significant. In contrast, for the non-SNI hind paw all scatter plots look random. Thus, FC between dorsal hippocampus and this specific part of somatosensory cortex specifically promotes neuropathic tactile allodynia. **Figure 4D, E** shows discovery and validation scatter plots, averaged across all four dorsal hippocampal seeds, for FC values from the dorsolateral thalamus (**fig. 4D**) and the limb region of the primary somatosensory cortex (**fig. 4E**). In both regions, discovery and validation indicate a positive correlation between ΔFC and ΔThreshold for the SNI hind paw, but no relationship for the non-SNI hind paw. Therefore, high FC values between dorsal hippocampus and the dorsolateral thalamus or the limb region of the primary somatosensory cortex are related with decreased neuropathic pain.

Overall, the chemo-fMRI whole-brain voxel-wise analysis shows that enhancing excitability of bilateral dorsal hippocampus modulates a complex network specifically involved in controllilng neuropathic tactile sensitivity. Using a strict discovery and validation approach we identify three brain regions most reliably related to tactile allodynia modulation by the dorsal hippocampal neuronal excitability.

### Large-scale network analysis identifies whole-brain and dorsal hippocampus functional connectivity related to neuronal excitability of the dorsal hippocampus

We analyzed the differential impact of PSEM^89s^, in contrast to saline, at a coarser scale, from the viewpoint of parceling the brain to 96 anatomically defined nodes and constructing resultant networks across these regions (**fig. 5**). Here, we used data from both scans 1 and 2, to enhance statistical power. The group averaged change in FC (between PSEM^89s^ and saline) is shown for all nodes, and for the 4 dorsal-hippocampus nodes (**fig. 5A, B)**. Increased and decreased FC for the full matrix and their strength distribution are also illustrated (**fig. 5C, D**). Overall, we observe twice as many connections increasing in strength than decreasing. Regarding the 4 dorsal-hippocampus nodes, increased connectivity is mainly with cortical targets, while decreases are with sub-cortical structures (see **supplementary table 3** for details). No obvious connectivity patterns differentiate between the 4 dorsal-hippocampus nodes. This network analysis results are in close correspondence with those observed for the seed-based analysis.

**Figure 5.**
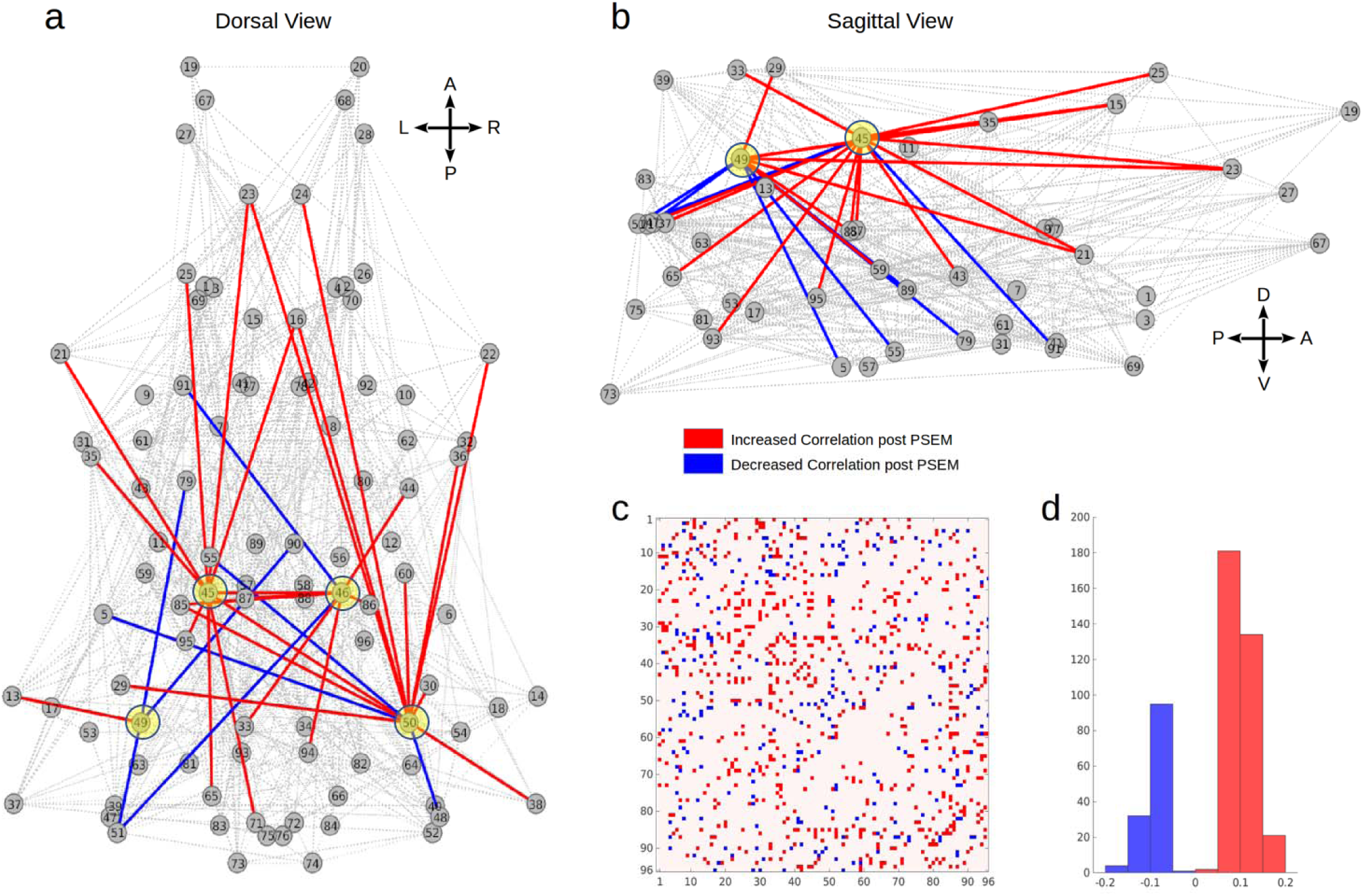
Functional network analysis, based on sub-dividing the brain into 96 clusters, identifies whole-brain and dorsal hippocampus connectivity changes with increased dorsal hippocampus neuronal excitability. **A, B**, Dorsal and lateral views of change in network connectivity, comparing PSEM^89S^ with the saline injection conditions. Change in connectivity across all nodes are shown in grey; increased and decreased connectivity for the 4 dorsal-hippocampus nodes (highlighted) are shown in red and blue, respectively. **C.** The full matrix is displayed, indicating increased and decreased connections between all 96 nodes. **D.** Histogram of the significant changes in the average correlation coefficient for PSEM^89S^ in contrast to saline. Contrasts were performed combining scan 1 and 2 data, using permutation testing, p <0.05 (see **supplementary table 3** for statistical details; brain regions corresponding to node numbers are in **supplementary table 4**). (A = anterior; P = posterior; L = left; R = right; V = ventral; D = dorsal).

### Anterograde tracing of hippocampal projections

The dorsal and ventral hippocampus showed non-overlapping projections to brain areas, many involved in fear extinction. Tracing was performed by injecting AAV8-mCherry into the dorsal (anteroposterior -3mm; mediolateral 1 mm; ventrodorsal 2.25mm) or ventral (anteroposterior - 1.8mm; mediolateral 2.25 mm; ventrodorsal 3mm) hippocampus, **figure 6, DH & VH**, respectively. The DH innervation of mPFC, BLA, and VH were scarce if any; whereas VH innervated all of these areas as well as DH. Projections in septum and mamillary complex were complementary: DH projected to the dorsal septum and VH to intermediate lateral and ventral septum; DH to supramammilary and VH to mammillary nuclei; DH to dorsal ACC and VH to ventral ACC. Both DH and VH projected to PAG, but VH more densely than DH.

**Figure 6.**
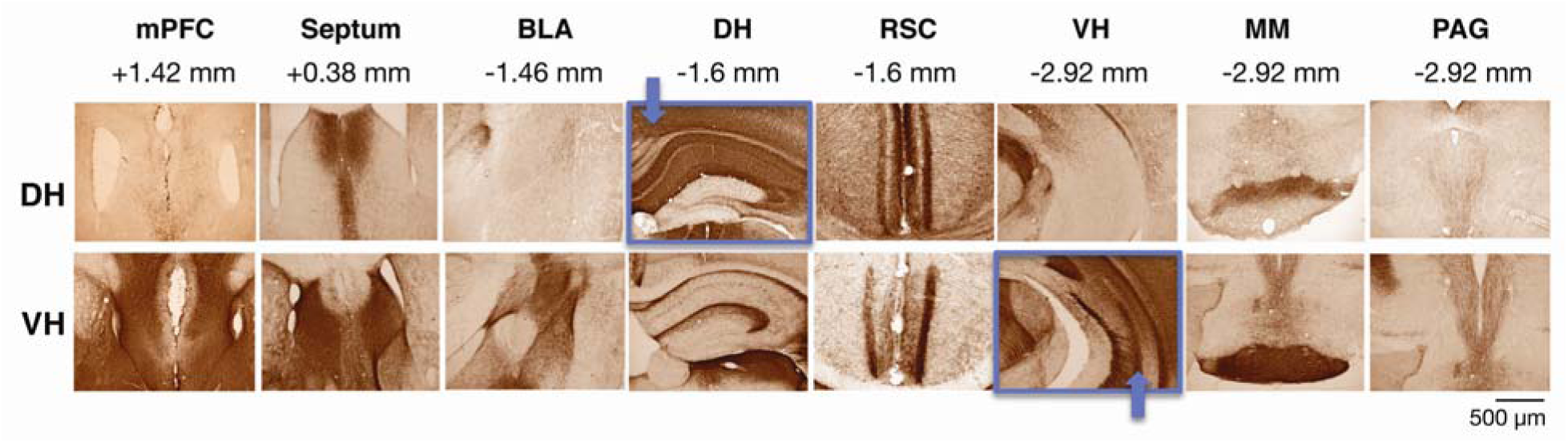
Projections from dorsal and ventral hippocampus. Tracing was performed by injecting AAV8-mCherry (UNC, 0.5µl/site) into the dorsal (anteroposterior -3mm; mediolateral 1 mm; ventrodorsal 2.25mm; blue arrow, box) or ventral (anteroposterior -1.8mm; mediolateral 2.25 mm; ventrodorsal 3mm; blue arrow, box) hippocampus. Immunostaining was performed using anti-mCherry antibodies (1:1,000, Abcam, ab167453) and visualized using diaminobenzidine. DH = terminations seen from dorsal hippocampus injection; VH = terminations from ventral hippocampus injection. mPFC = medial prefrontal cortex; BLA = basolateral amygdala; RSC = retrosplenial cortex; MM = mammillary bodies; PAG = periaqueductal grey.

## Discussion

We uncovered that increasing excitability of the dorsal hippocampus, either by glutamate microinjection or by chemogenetics, could alleviate neuropathic pain behaviors. Both manipulations reversed neuropathic behaviors, and glutamate injection in the dorsal hippocampus diminished exaggerated tactile and thermal behaviors, in two different models of neuropathic pain. Unilateral glutamate injections reversed neuropathic signs bilaterally when both hind limbs exhibited hyper-sensitivity to touch (C-SNI); while in models where the injury resulted only in ipsilateral hypersensitivity (L5 SNL, U-SNI), ipsilateral (glutamate) or bilateral (chemogenetic) increased excitability of the dorsal hippocampus only diminished responsiveness for the hind paw/paws exhibiting neuropathic hyper-sensitivity. Both manipulations diminished neuropathic signs transiently (1-2 days with glutamate injections, about 1 hour with chemogenetics), and with glutamate we could also demonstrate a dose-response effect, and complete blockade of efficacy when the glutamate injection was preceded by injection of an NMDA receptor antagonist. Efficacy of reversal of neuropathic signs, with both manipulations, was observed repeatedly at various time delays from peripheral injury (tested from 4 to 20 days from induction of peripheral injury), implying engagement of the dorsal hippocampus in neuropathic pain soon after injury and persistently. The fact that the 5HT3 receptor activation by PSEM^89s^ (but not with saline), transiently, repeatedly and specifically reversed tactile allodynia provides direct evidence regarding the role of dorsal hippocampal neurons in controlling neuropathic pain. Therefore, we can conclude that, at least in these rodent models of chronic neuropathic pain, the state of excitability of the dorsal hippocampus, but not the ventral hippocampus, specifically and causally controls chronic neuropathic behaviors.

We next used chemo-fMRI in minimally restrained, trained and awake rats, to identify the brain circuitry underlying dorsal hippocampal control of neuropathic pain. Using a discovery and validation approach, we identified the brain circuitry activated/deactivated during increased excitability of the dorsal hippocampus specifically related with diminution in neuropathic pain. Extended, cortical and sub-cortical circuits were identified. Yet only a select brain regional FC changes could be validated, and we consider these as the most robust FC outcomes related to tactile allodynia. Our tracing study indicated that dorsal hippocampus projections are more restricted and non-overlapping with ventral hippocampal projections.

Our obtained results are generally consistent with older observations: A human PET study shows bilateral hippocampal deactivation in patients with mono-neuropathy [34]; while rodent studies show persistent decreases in excitability of dorsal hippocampus CA1 pyramidal neurons in the formalin model (a mixed nociceptive and neuropathic model for persistent pain that only lasts for a few days) [24], and electrical stimulation of the dorsal hippocampus is reported to lead to anti-nociceptive behaviors [36]. Abnormal hippocampal glutamatergic activity has also been implicated in the pathophysiology of chroic pain [54], and decreased hippocampal LTP is observed in slices from neuropathic pain rats [26]. After chronic constriction injury of the sciatic nerve, hippocampal CA1 glutamate concentration is decreased, whereas GABA concentration is increased and correlated with pain thresholds [41]. There is also evidence for inhibition of GABAergic systems of the dorsal hippocampus by microinjection of bicuculline increasing antinociception, while muscimol promoting pronociception, suggesting blockade of GABAergic neurons activity may be antinociceptive [20]. Thus, increasing the local concentration of glutamate in dorsal hippocampus may inhibit mechanical allodynia by rebalancing local excitatory/inhibitory neurotransmitters. NMDARs can activate the ERK/CREB signaling pathway [8]. Our previous work has shown that ERK expression is decreased in the hippocampus after SNI; while here we found the inhibitory effect of glutamate on mechanical allodynia was blocked by an NMDA antagonist. ERK/CREB pathway can be activated by increases in intracellular calcium level through NMDA receptors. Thus it is possible that glutamate injected into the dorsal hippocampus increased ERK activation by binding to NMDA receptor. Overall, then we suggest that glutamate microinjection effects were mediated mainly by increasing excitatory local circuits, resulting in activating and deactivating the circuitry we identify with chemo-fMRI. Still, it remains unclear whether increased net excitability or simply disruption of local activity patterns underlie the overall observed results. The latter may be sufficient to relieve neuropathic pain either by perturbing memory retrieval and storage processes by interfering with coordinated information flow within the intra- and inter-regional cirtcuits (for example, by disrupting local field potentials [29]); and/or by enhancing memory extinction processes [47] as we have shown that contextual fear extinction is inhibited in neuropathic pain [32] and distinct dorsal hippocampus neurons and molecular processes underlie extinction of contextual fear [21; 22; 42; 48]. Parameters that control the duration of the efficacy of glutamate on neuropathic pain, in a dose dependent manner, remain unclear, and may require disentangling between storage and extinction circuitry specifically in relation to chronic pain.

As the chemogenetic study was done in rats where specific molecular tags for neuronal types were not available (cre independent virus was expressed), our approach infected all neurons within the virus injection site. Thus, the specific neuronal populations within the dorsal hippocampus controlling neuropathic behaviors remain unclear, and future experiments will be required to clarify this point. However, given the effects of PSEM^89S^ and glutamate replicated each other, the effect of glutamate could be blocked by NMDA receptor antagonist, NMDA plays an excitatory role on pyramidal neurons, and formalin injection in the hind paw suppresses CA1 pyramidal cell in dorsal hippocampus by exciting GABAergic and suppressing pyramidal activity [24], we surmise that the net effect of PSEM^89S^ mediated unmasking of 5HT3 channels is an increase in pyramidal neuronal activity, the efferent projections of which then modulated behavior. Still we cannot rule out that the chemogenetic effects are also due purely to disruption of local activity patterns. It is striking that a subtle manipulation of membrane properties of dorsal hippocampal neurons results in a robust and reproducible alleviation of tactile allodynia in every animal tested. We intentionally designed the chemogenetic portion of the study to infect a large portion of the region (bilaterally, encompassing anterior and posterior portions of dorsal hippocampus) to insure observing some behavioral changes. Yet, in all animals tested, this procedure was efficacious.

The chemo-fMRI study involved a limited number of animals. To compensate, we restricted statistical analyses, and used a discovery/validation approach. In the voxel-wise analysis we only performed a single statistical contrast (correlation of ΔFC with ΔThreshold being > 0, or < 0); in the coarse-grained network analysis we used permutation testing for the nodes of interest. In the behavioral correlation analysis, we only report regions with corrected outcomes that pass a p-value threshold > 0.0001, and of those we emphasize the 3 regions that passed validation. The two network analyses results show good similarity; both indicate increased connectivity to multiple cortices and decreased connectivity to sub-cortical regions. Lateralization of this circuitry remains unclear, although our behavioral results suggest bilateral influences. The connectivity analysis for the four hippocampal subdivisions does not show any clear specificity, but the issue requires more focused future studies. Given the projections of the dorsal hippocampus to the thalamus [11; 50] and cortex [11; 46], which are consistent with the tracing results described here, we presume that the thalamic and retrosplenial functional connectivity identified here are mainly monosynaptic in nature; while the somatosensory, motor, insular and prefrontal connections, as well as the sub-cortical connections are more likely polysynaptic networks. Three dorsal hippocampus connected networks, sub-regions of the primary somatosensory cortex and the dorsolateral thalamus, survived validation and thus are considered most reliable circuitry underlying control of neuropathic behavior. We suspect additional networks underlie the behavioral modulation but these await future larger studies.

Given the dorsal hippocampus is preferentially engaged in episodic memories and spatial navigation functions, whereas the ventral hippocampus implicated in stress, affect and emotional memories [1; 19; 39; 45], in agreement with earlier observations [3] our results highlight the independence of episodic memory circuits from negative mood circuits in chronic pain, and extends this notion by demonstrating the former are tightly related to neuropathic pain behaviors. Complementary results have been shown recently in yet another rat neuropathic pain model (partial sciatic nerve ligation), where treatment with D-serine could ameliorate anxiety-like behaviors but not tactile allodynia [54], presumably through ventral, rather than dorsal, hippocampal circuits. It is noteworthy that in the current study increased, or perturbed, excitability of a large portion of the dorsal hippocampus resulted in local intrinsic FC and changes in FC with multiple regions outside of the hippocampus, but we did not observe spread of FC to the ventral hippocampus, consistent with our tracing data showing lack of projections from dorsal to ventral hippocampus. The latter is again consistent with independence of the two regions from each other. Still, the differential reorganization of the dorsal and ventral hippocampi with transition to chronic pain remains unclear and needs additional studies.

Our results show the power of combining chemogenetics with fMRI, especially in awake animals, where behavior can be directly linked to specific brain circuitry. The methodology unravels brain circuitry for excitability of a specific group of neurons, linking them to chronic pain behavior. To our knowledge this is the first application of the chemo-fMRI technique, although a recent proof of concept report shows feasibility of a similar approach where Designer Receptors Exclusively Activated by Designer Drugs (DREADD)-technology combined with pharmacological-fMRI in anesthetized rats [40]. Our approach links whole-brain FC changes that could be directly compared to similar analyses in humans, potentially providing evidence for model equivalence across species, lending novel mechanistic insights to human data. We have reported in the past on hippocampal FC changes in humans with chronic back pain, and in subacute back pain during the transition to chronic pain [31], as well as optogenetic results in mice with SNI regarding hippocampal efferents to nucleus accumbens shell D2 iSPNs [37]. All three studies show similarities and complimentary information. However, we should note that there are important methodological differences (the human study entailed whole hippocampus FC analyses, during active tasks; the mouse study examined AMPA/NMDA ratio for ventral hippocampal inputs to D2 iSPNs; the current rat study examined dorsal hippocampus FC when excitability was transiently chemogenetically enhanced). In all three studies (3 species), we observe either FC, or physiologically identified, changes in hippocampal outputs. In the human study, transition to chronic pain is accompanied with increased FC intrinsically, and extrinsically from the anterior hippocampus to multiple cortical targets; a close parallel and complimentary to the current rat results, which enables the conclusion that the human FC changes are also most likely causally controlling extent of chronic pain perception. All three studies together suggest the conclusion that transition to chronic pain is likely accompanied with decreased dorsal and increased ventral hippocampal excitability, and FC (monosynaptic or polysynaptic) distinct changes contributing to both pain perception and to the negative affective properties of chronic pain.

## Conceptual Implications

The overall conceptual implication of the current study is the notion that disrupting dorsal hippocampal, episodic/cognitive memory-related retrieval or extinction processes, information flow to thalamic and cortical targets reverses behavioral signs of neuropathic pain. As these circuits are providing a “map” for sensorimotor planning and behavioral execution [58], based on accumulated contextual and experiential memories, renormalization of pain behavior with excitation/disruption of these circuits suggests that the acquired memories (with persistence of pain), coupled with inability to extinguish, is directly involved in chronic pain behavior/perception. Hippocampal “map”-s reflect locations in spatially organized environments and also map moments in temporally organized experiences, and the integration between these maps is thought to reflect a generalized mechanism for organizing memories [17]. Therefore, these maps must undergo dramatic reorganization/suppression with neuropathic pain, the details of which remain entirely unknown.

Our results show dorsal hippocampal control over neuropathic pain conditions soon after the injury and throughout the window where we conducted these studies (from 4 to 20 days after injury). There is ample evidence that, within this time window, both peripheral and spinal cord nociceptive circuitry are sensitized [57]. Moreover, the behavioral measures we have used in this study assess primarily spinal cord mediated reflexive outcomes, which are a reflection of the extent of spinal cord nociceptive sensitization. Thus, our results establish the first evidence for a potent control by dorsal hippocampus-thalamus/somatosensory cortex circuitry on spinal cord nociceptive circuitry, causally modulating the level of this sensitization by excitability of dorsal hippocampal circuits.

Ten years ago, Apkarian [2] advanced the theoretical concept of chronic pain being a state of extinction-resistant emotional learning. A corollary to the concept is the continuous acquisition and consolidation of memories through emotional learning, accompanied with minimal opportunity for extinction processes to overcome or diminish the emotional impact of the condition. There is ample evidence that emotional states influence learning and memory [56]. The present results provide the most direct evidence that the negative emotional state of persistence of neuropathic pain gives rise to the accumulation of long term pain-related extinction-resistant memories, which control neuropathic behaviors (and most likely human perception and behavior, given the close parallels seen in human hippocampus FC with transition to chronic pain [31]). Stress, as an alternative example for a negative emotional state, is shown to facilitate and/or impair both learning and memory, depending on intensity and duration [23; 51; 52]. The present results complimentarily suggest that persistent/neuropathic pain is itself, to a large extent, the accumulation of memories related to the experience of pain. Emotional learning engages large components of the brain limbic circuitry, and evidence suggests circuit specificity: the amygdala modulating memory consolidation; the prefrontal cortex mediating memory encoding and formation; the ventral basal ganglia providing motivational drive for learning and generalization; and the hippocampus mediating learning and memory retention [53; 56]. Animal studies, besides the evidence provided here for the dorsal hippocampus, now indicate causal relationships for large parts of this circuitry in neuropathic or persisting pain conditions, especially regarding the medial prefrontal cortex, nucleus accumbens, and amygdala; and parallel evidence continues to accumulate in human chronic pain studies as well (see review [4]). Thus, there is compelling evidence that brain circuits causally controlling neuropathic pain are much more extended in the brain than previously suspected. How all these systems are interrelated and in turn interact with nociceptive representation in the cortex and at the level of the spinal cord remains a critical challenge.

## Materials and Methods

### Animals

Sixty-seven adult male Sprague Dawley rats weighing 200-250 g were used in these experiments. The animals were group housed and given free access to standard rodent chow and water. Some of the described experiments were done in Sun Yat-sen University, Guangzhou, China, while others were done in Northwestern University, Chicago, USA.

The room was kept at 21 ±2°C temperature and 30-60% humidity, under a 12/12 h light/dark cycle. Handling and testing were performed during the light period. To minimize stress, they were handled regularly before operation and before behavioral testing. All studies were approved by the Animal Care and Use Committee of Northwestern University or by the Local Animal Care Committee (Sun Yat-sen University) and with the ethical guidelines for investigation of experimental pain in conscious animals [59]. Totally 10 rats were excluded: of studies done in China, 1 died after surgery and 7 animals did not develop bilateral neuropathic behavior; in studies done in the USA, 2 animals’ head-posts were dislodged after implantation. No randomization was used for assigning the animals to different treatment groups.

### Neuropathic pain models

#### Spared nerve injury (SNI)

Spared nerve injury was carried out following the original description [16]. Briefly, rats were anesthetized with isoflurane (1.5-2.5%) and a mixture of 30% N_2_O and 70% O_2_. The sciatic nerve of the left leg was exposed at the trifurcation of peroneal, tibial, and sural branches. The common peroneal and tibial nerves were ligated and cut, whereas the sural nerve was left intact. For the animals in the sham group, the sciatic nerve was only exposed but not disturbed.

#### Spinal nerve ligation (SNL)

Lumbar segment 5 spinal nerve ligation was done following the original description [25]. Briefly, after rats were anesthetized with isoflurane (1.5-2.5%) and a mixture of 30% N_2_O and 70% O_2_, the L5 transverse process was removed to expose the L5 spinal nerve. L5 spinal nerve was then isolated carefully and ligated tightly with 6 – 0 silk thread 5-10 mm distal to the L5 DRG.

#### Drug administration with brain microinjection

L-Glutamate acid, monosodium salt monohydrate and D(−)-2-Amino-5-phosphonopentanoic acid (AP-5) used in this study was purchased from Sigma (ST, Louis, MO, USA). For the brain microinjections, rats were anesthetized with sodium pentobarbital (50 mg/kg, i.p.) and permanent guide cannulas (23 gauge) were implanted in the dorsal or ventral hippocampus, ipsilaterally to the peripheral injury. The stereotaxic coordinates were: AP -3.5 mm; ML -2.0 mm; DV -3.2 mm (dorsal hippocampus), and AP -4.8 mm; ML 4.6 mm; D -8.1 mm (ventral hippocampus), [33]. The cannula was secured by dental cement and anchored to stainless steel screws fixed to the skull. To prevent clogging and infection, the stainless-steel obturator was inserted into the guide cannula. The animals were allowed to recover for six days after surgery. Infusion was performed with Hamilton syringe connected to a 30-gauge injector. Drugs were infused (1µl over 1 minute) into the dorsal or ventral hippocampus, ipsilaterally relative to the peripheral injury. The injection needle was kept in place for 60 s to allow the drug to completely diffuse from the tip, and then the obturator was reinserted into the guide cannula. AP-5 was injected through the cannula placed in the dorsal hippocampus, 30 min before glutamate injection.

### Assessment of mechanical allodynia

All behavioral tests were done in a blinded paradigm.

1. Mechanical response thresholds of the animals before and after SNI was assessed with the up-down method [14]. Briefly, animals were placed in a Plexiglass box with a wire grid floor and allowed to habituate to the environment for 10-15 minutes. Filaments of varying forces (Stoelting Co, USA) were applied to the lateral plantar surface of the hind paw. Filaments were applied in either ascending or descending strengths to determine the filament strength closest to the hind paw withdrawal threshold. Each filament was applied for a maximum of 2 seconds at each trial; paw withdrawal during the stimulation was considered a positive response. Given the response pattern and the force of the final filament, 50% paw withdrawal threshold was calculated following the method [14].

2. To test for mechanical response sensitivity, the incidence of paw withdrawal to four different forces of filaments (2, 6, 15, and 26 g) was measured according to previous work [18; 55]. Each filament was applied once a second to the plantar surface for eight times. Ten trials were performed on each hind paw. The percentage of positive trials was recorded.

### In vivo chemogenetic activation of hippocampus

Adult (8–10 weeks; 200–250 g) male Sprague Dawley rats were group-housed until surgery. As previously described [37], stereotaxic-guided virus injection surgeries were performed under isoflurane anesthesia and using a stereotaxic instrument (Kopf Instruments, Tujunga, CA). Four small holes (two per side) were drilled through the skull and bilateral virus injections were made using glass injection pipette (Drummond Scientific Company). To activate dorsal hippocampus, AAV9-SYN-PSAM-L141F-Y115F-5HT3HC-GFP (titer: 2.19× 10^13^ genomes/ml, Virovek, Inc.) were injected into bilateral dorsal hippocampus (bregma -3 mm; lateral ±1.7 mm; ventral -3.3 mm for rostral coordinates; bregma -4.2 mm; lateral ±2.7; ventral -3.3 mm for caudal coordinates; 50 nl / side). Head-posts were implanted 2-3 weeks after virus injection. Following the 2-week acclimation period, SNI surgeries were performed and 4-5 days after SNI surgery, the paw withdrawal threshold was assessed at pre- and 0.5-2 hours post i.p. injection of either saline or PSEM^89s^ (30 mg/kg). The virus injection sites were verified by immunohistochemistry.

## Electrophysiology

Sagittal brain slices containing dorsal hippocampus were obtained from PSAM virus injected rats. Experiments were performed as previously described (Ren et al., 2016). Briefly, the rats were anesthetized with ketamine/xylazine and perfused transcardially with ice-cold artificial CSF (ACSF). Brains were rapidly removed and sliced (VT1200S vibratome, Leica Microsystems), and then the slices were incubated and transferred into recording chamber. Using patch clamp recording, The GFP positive pyramidal neurons in dorsal hippocampus slice were recorded under current clamp recording mode. PSEM^89S^ was applied to the bath using a by gravity-driven drug application system. Picrotoxin (100 μM) was added to block GABA_A_ receptor-mediated inhibition, and a combination of CNQX (10 μM) and D-AP5 (50 μM) were used to block fast glutamatergic transmission.

## Immunohistochemistry

Rats were anesthetized with ketamine/xylazine and transcardially perfused with 4% formaldehyde in 0.1 M phosphate buffer, pH 7.4. After post-fixed for 24 hours in 4% formaldehyde, the brains were cryoprotected in 30% sucrose in PBS. Cryoprotected brains were sectioned serially at 30 μm in sagittal or coronal planes on a freezing-stage microtome. All of the sections were blocked with 2% goat serum in 0.1% Triton X-100 for 1 h at room temperature and incubated over two nights at 4 °C with primary antibodies for green fluorescent protein (GFP, 1:3000, Millipore). The immunoreactions were either visualized by secondary antibodies coupled to Alexa 488 (Invitrogen) or detected by staining with 3, 3′-diaminobenzidine (DAB). For all cases, representative pictures of selected regions in the GFP-positive sections were collected to ascertain the viral injections properly targeted the dorsal hippocampus.

## Awake resting state functional MRI (rsfMRI)

These experiments closely follow methods we developed and described recently [12; 13].

### Head-post implantation

Surgery for head-post implantation was performed under anesthesia. Rats were initially anesthetized with 3.5% isoflurane mixed with 30% N2 and 70% O2, then transferred to a stereotaxic device and mounted using blunt ear bars that do not break the eardrums. Anesthesia was continued at a lower concentration sufficient to block motor responses to pinching the hind limbs. The head fur was shaved, and eyes were covered with ointment to prevent drying out and corneal infection. After disinfecting the skin overlying the skull, the scalp was cut longitudinally and the skin retracted from the cranium. The head-post was placed at the midpoint of bregma and lambda, and fixed to the skull with dental cement. The skin wound was treated with antibiotics. The animals were then released from the stereotaxic frame and kept warm using a heat lamp. After head-post implantation, rats were given at least two weeks of rest to recover from the surgical preparation before the start of acclimation procedures.

### Acclimation procedure

For animals that underwent awake scanning, they first received a surgical implantation of a head post to aid in stabilization during scanning. They were then subject to an acclimation training period in which rodents were first introduced to snuggle sacks until they began to voluntarily walk into the snuggle sacks. Subsequently, snuggle sacks were adjusted around the rodents for a snug fit. Once acclimated to their snuggle sacks, the rodents were transferred to cradles and then, the animal’s body was secured to cradles with Velcro straps. The head plate was affixed to the head and subsequently secured to sidewalls mounted on the cradle. Rodents were then acclimated to head restraint with a graded training procedure. With repeat of acclimation, the rodent’s head was gradually held firmly to the head-plate that was securely mounted on the cradle. To reduce the stress induced by MRI experimental environment, rodents were exposed to digital recording of the sounds generated by gradient switching in the magnet during fMRI so they were familiar with loud and sudden noise that mimics a typical MRI session.

Rats were habituated to the body restrained system and the MRI environment in 30-min sessions/day for 8–10 days, within a 2-week period. The animals were first acclimated in the mock scanner box that simulated the MRI scanner environment and, during later times, directly in the MRI scanner. Rodents were rewarded with treats at the end of each session. The training was performed after the head-post implantation and before the SNI surgery and it was repeated after surgery prior the collection of the imaging data.

### Awake Functional magnetic resonance imaging

Each animal underwent a sequence of three imaging scans assessed the same day at baseline –no injection-, 1 hour after or saline systemic injection, at day 4 or 5 after neuropathic surgery. The order of the saline and PSEM injections were randomized between animals and at least 2 hours period was left in between both to exclude any remaining effect of the drug. A total of 9 rats were used in the experiment. Animals whose head-post became detached during retraining sessions were not imaged. In addition, data from one rat at baseline session with excessive motion artifacts was likewise discarded. Thus, we retained scans for 7 SNI animals after 1 hour of PSEM and saline, and 6 animals at baseline.

### Equipment for awake rodent fMRI

The instrument -including MRI holder, cradle, head plate, head-post and mock scanner box-was designed using SolidWorks software and was three-dimensional printed (ProofX, IL) using a semitransparent, medical grade material (PolyJet photopolymer MED610, Stratasys) that we had previously tested for magnetic susceptibility. Screws (Small Parts) and fastener were made with Ultem PEI polyetherimide. Custom-made “snuggle sacks” were designed to tailor for rodents weighting 300–400 g. Snuggle sacks included openings that provided access to the head-post. More details can be found in [13].

### Magnetic resonance imaging acquisition

All magnetic resonance experiments were performed on a Bruker 7T Clinscan horizontal magnet. A 2-channel volume resonator was used for radio frequency transmission, and a 2-cm-diameter surface coil was used for signal detection. Blood oxygenation level-dependent (BOLD) contrast-sensitive T2*-weighted echo-planar imaging (EPI) was acquired for functional images with the following parameters: gradient-echo, 26 oblique transverse slices, repetition time 2007.5 milliseconds, echo time (TE) 18 milliseconds, in-plane resolution 0.348 mm, slice thickness 0.5 mm, number of repetition 240. Additional images were collected to optimize registration, including a T2-weighted anatomical image, having identical spatial dimension as in the functional images. EKG and body temperature were monitored were monitored during all fMRI experiments.

### Image preprocessing

Functional MRI data was preprocessed using FSL 5.1 FEAT (FMRIB’s Software Library, http://www.fmrib.ox.ac.uk/fsl) default settings, and included motion correction with MCFLIRT, interleaved slice timing correction, skull removal, and spatial smoothing (FWHM) with a Gaussian kernel of 0.7 mm (not default), chosen because it is roughly 1.5 times the largest dimension of the voxel. BOLD time courses were forward and backward bandpass filtered at 0.01 to 0.08 Hz with a 4th order butterworth. Functional images were then registered using FLIRT, first to the individual anatomical T2 image with six degrees of freedom (not default) then this was registered linearly to the standard brain using twelve degrees of freedom. We used an in-house manually drawn template to identify cerebrospinal fluid and white matter voxels in a standard brain image. Average time courses from all the white matter (WM) and cerebrospinal fluid (CSF) as well as those pertaining to global whole brain signal were extracted using the FSL fslmeants function. CSF, WM, global signals and six motion parameters (3 translation and 3 rotation) were regressed out from the images using FSL’s fsl regfilt linear regression function.

### Seed-based connectivity

Functional connectivity (FC) maps were generated in Matlab for the 4 seeds where the PSAM virus was injected. Using functional images registered to standard space, the average BOLD time series from 3-voxel radius spheres surrounding the virus injection coordinates were extracted, and individually correlated against all voxels in the brain using Pearson’s correlation. Average FC maps across all 4 seeds were then contrasted between PSEM and saline injection conditions (paired within animal contrasts). A higher-level analysis was then performed to identify FC changes that correlated with the behavioral differences, observed for the injured paw, between PSEM and saline conditions for results collected from the SNI hind paw. Individual animal FC difference maps (between PSEM and saline) were correlated with change in tactile allodynia between PSEM and saline. FSL FLAMEO was again used to identify z-stat maps, corrected for multiple comparisons using intensity z>2.3 and cluster p<0.05 thresholds.

The above analysis was the discovery phase, which was based on one set of fMRI scans collected in each animal, at 1-hour after PSEM or saline injections. We had also collected a second fMRI scan immediately after termination of the first, and this scan was reserved for a validation analysis. Preprocessing and post-processing (to generate FC maps for each seed) steps for the validation data were identical to those we used in the discovery data. However, resultant maps were not thresholded, only region of interest FC values were extracted and studied. The validation FC maps, reflecting individual animal, seed, and condition specific outcomes were used to extract FC values at the coordinate of interest derived from the discovery results. Resultant regions were considered to be validly reflecting tactile changes only if the joint p-value, for observed correlations between discovery and validation, was significant at p<0.0025 (0.05*0.05). Only 3 (of 8 tested; for highest probability 4 positive and 4 negative FC changes) regions survived this criterion.

In the virus injected animals, we collected one more resting state fMRI immediately after determining tactile thresholds 4-5 days after SNI, but just prior to saline or PSEM injections. We used the same preprocessing and post-processing steps for these scans as well. These data were only used to extract mean FC map, across the 4 seeds.

### Functional networks

We also constructed FC network for the whole brain, to examine its properties between PSEM and saline conditions, especially for the nodes where PSAM was introduced. The procedure closely follows our earlier methodology [5]. Briefly, the brains were segmented to 96 regions (ROIs, 48 per hemisphere) using a standard rat atlas [43]. The time course of BOLD signal for all voxels in each ROI were averaged. Resultant average BOLD signals were used to calculate Pearson correlation coefficients between pairs of ROIs; resulting in a 96×96 correlation matrix per scan. Data from scan 1 and 2, for PSEM^89s^ and saline injections (14 scans per condition) were used to determine functional connectivity changes between PSEM^89s^ and saline conditions. We next calculated average correlation matrices of each condition, and the absolute difference between the PSEM and saline averages. In order to evaluate the statistical significance of these changes, we employed a permutation test, where we randomized the labels of the correlation matrices, and calculated the difference between the new groupings. We repeated the randomization 50,000 time to obtain the Null Hypothesis Distribution (NHD) for each pairwise connection – the distribution of probable changes in the strength of a connection when the groups are not necessarily different. We then used the NHDs to select the connections with a statistically significant change (one-tailed p-value < 0.05).

### Tracing hippocampal projections

Nine week old male C57BL/CN mice obtained from a commercial supplier (Harlan) were used for these experiments. Mice were individually housed and allowed ad libitum access to food and water. Tracing was performed by injecting AAV8-mCherry (UNC, 0.5µl/site) into the dorsal (anteroposterior -3mm; mediolateral 1 mm; ventrodorsal 2.25mm) or ventral (anteroposterior - 1.8mm; mediolateral 2.25 mm; ventrodorsal 3mm) hippocampus. Immunostaining was performed using anti-mCherry antibodies (1:1,000, Abcam, ab167453) and visualized using diamino benzidine as described previously [15].

### Quantification and statistics

Data were expressed as means ± SEM, or as median and quartiles (for small number of observations). One-way or two-way analysis of variance with repeated measures followed by Bonferroni’s post hoc test was performed to analyze the behavioral outcomes. The criterion of statistical significance was P < 0.05. Although no power analysis was performed, the sample size was determined according to peers’ and our previous publications in behavioral and pertinent molecular studies. No randomization was used for data collection, but all initial analyses were performed blinded to the condition of the experiments. All brain imaging maps were corrected for multiple comparisons, and the final regions of interest were limited to ones that survived our rigorous discovery and validation procedure.

